## Supplemental Figures and Movie Legends for "Stepwise transmigration cascade of T and B cells through the perivascular channel in lymph node high endothelial venules"

**Contents:**

Figures S1-S12

Movies S1-S9

Fig. S1.

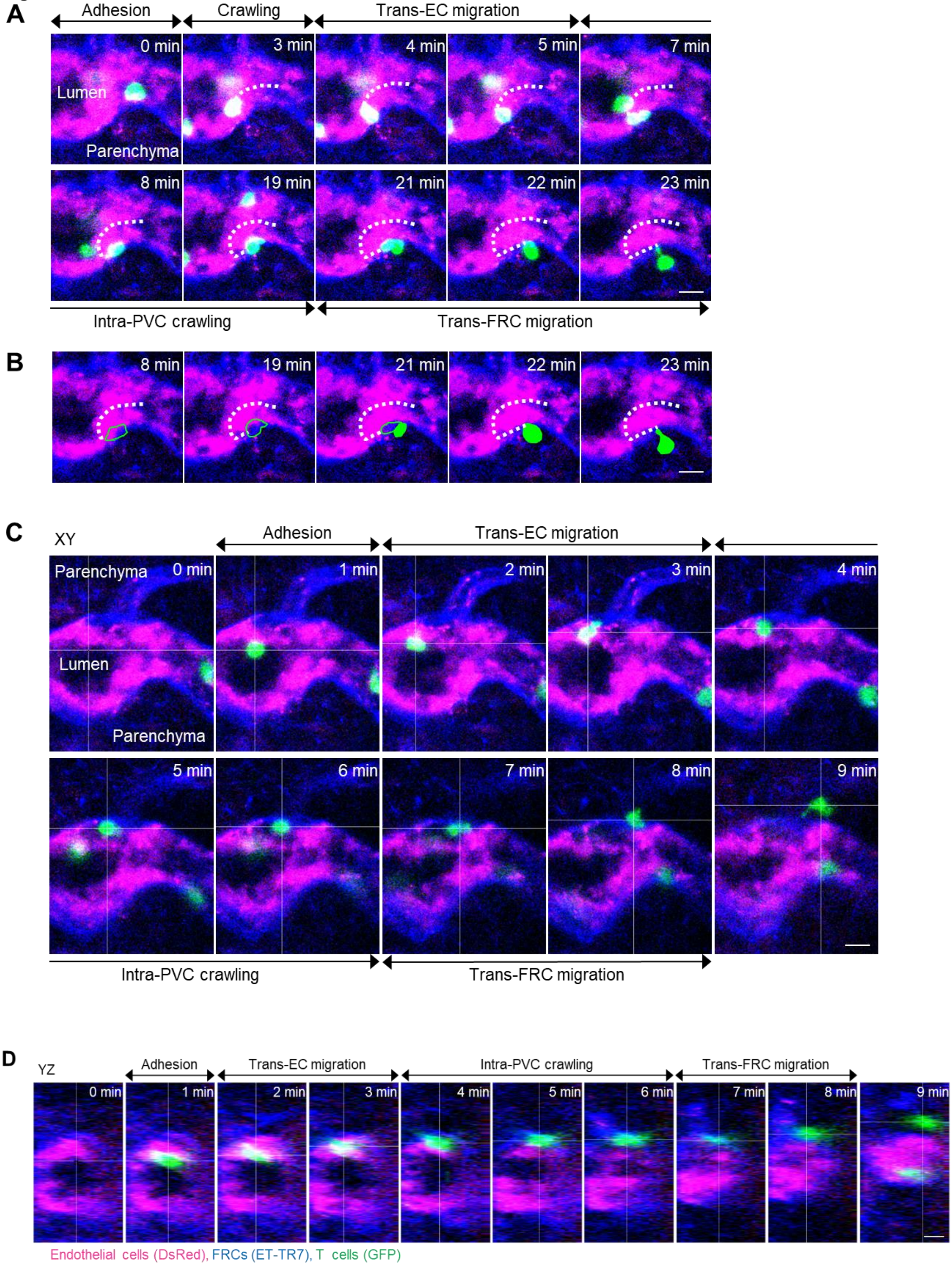

A
Adhesion
Crawling
Trans-EC migration
0 min
3 min
4 min
5 min
7 min
Lumen
Parenchyma
8 min
19 min
21 min
22 min
23 min
Intra-PVC crawling
Trans-FRC migration
B
8 min
19 min
21 min
22 min
23 min
C
XY
Adhesion
Trans-EC migration
Parenchyma
0 min
1 min
2 min
3 min
4 min
Lumen
Parenchyma
5 min
6 min
7 min
8 min
9 min
Intra-PVC crawling
Trans-FRC migration
D
YZ
Adhesion
Trans-EC migration
Intra-PVC crawling
Trans-FRC migration
0 min
1 min
2 min
3 min
4 min
5 min
6 min
7 min
8 min
9 min
Endothelial cells (DsRed), FRCs (ET-TR7), T cells (GFP)

**E**

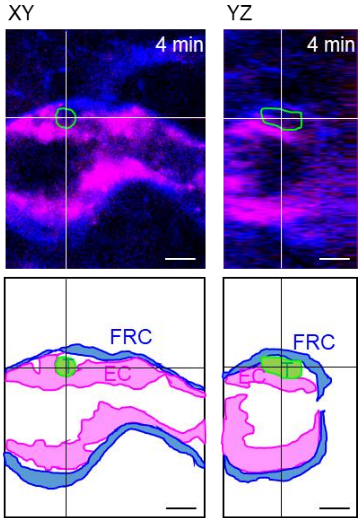


**Fig. S1.** (**A-B**) Serial single z-frames of Fig.1C showing the stepwise migration process of a T cell across an HEV; adhesion to EC, intraluminal crawling, trans-EC migration, intra-PVC crawling, and trans-FRC migration. The dotted line indicates the T cell track. (**B**) The intra-PVC segments of the T cell in green outline. (**C-D**) Additional example of the stepwise migration process of a T cell across the HEV. The image planes were selected based on the center of the T cell. (**C**) Serial single z-frames (XY plane) and (**D**) serial YZ-cross sections. (**E**) Images and illustrations corresponding to 4 min in Fig. S1C-D. The T cell is between endothelium and FRCs. The intra-PVC segments of the T cell in green outline. Scale bars, 10 μm.

**Fig. S2.**

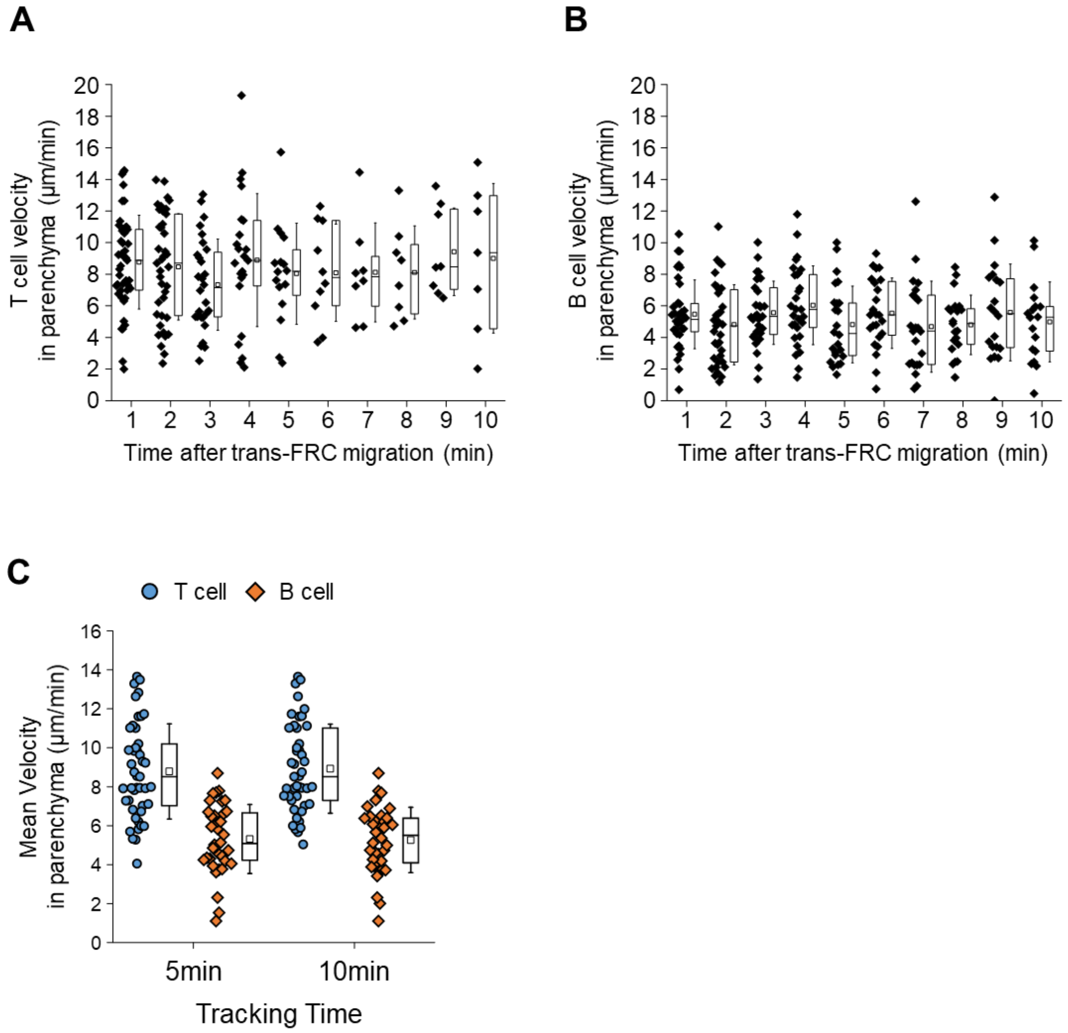


**Fig. S2.** (**A-B**) There is no significant change in T and B cell instantaneous velocities for 10 minutes after trans-FRC migration (One-way ANOVA Tukey's test). (**C**) Mean velocities inside parenchyma for 5 and 10 minutes are not significant different. Each symbol represents a single cell. The box graph indicates 25th and 75th percentiles; the middle line and whiskers of the box indicate the median and standard deviation, respectively; the small box represents the mean value. Four and 3 mice were used for the analysis of T and B cells, respectively.

**Fig. S3.**

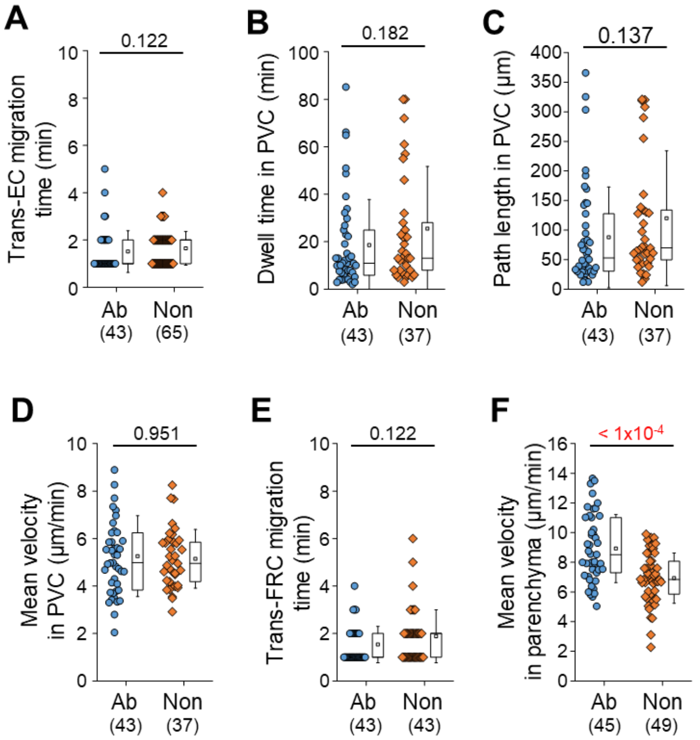


**Fig. S3. Effect of footpad injection of anti-ER-TR7 antibody on T cell transmigration across HEV.**

(**A-E**) There is no significant difference between antibody-injected group (Ab) and non-injected group (Non) in T cell migration from trans-EC migration to trans-FRC migration. Non-injected means that no substance is injected into a footpad of mouse. (**F**) The injection of anti-ER-TR7 antibody (10 μg/50μl) increase the mean velocity of T cells inside parenchyma within 10 min after the trans-FRC migration. Each symbol represents one single. The box graph indicates 25th and 75th percentiles; the middle line and whiskers of the box indicate the median value and standard deviations, respectively; the small box represents the mean value. The number of analysed cells is indicated below the graph. Four and 2 mice were used for the analysis of antibody treated and non-treated groups, respectively. P values were calculated with the Mann Whitney test.

**Fig. S4.**

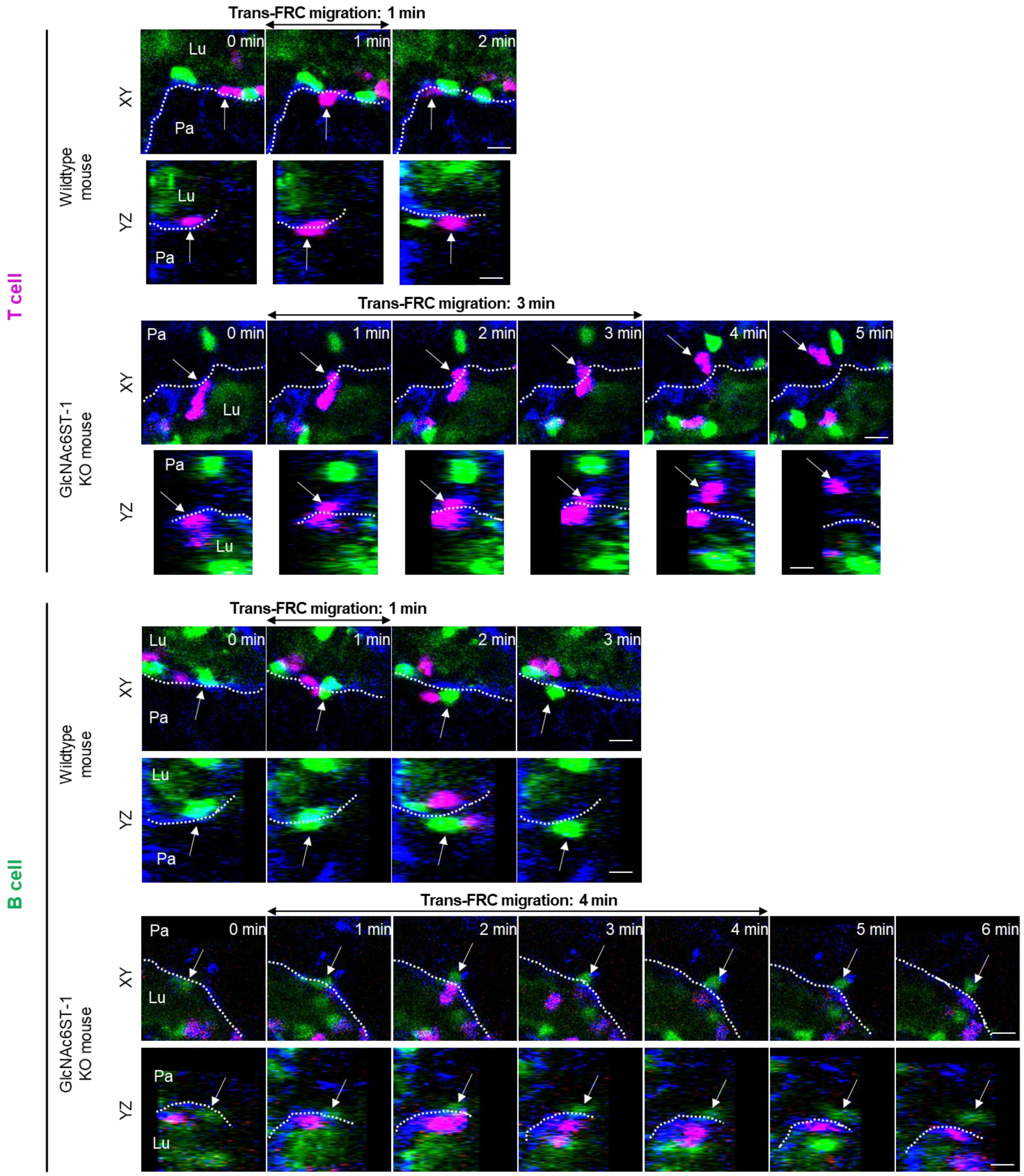


**Fig. S4.** Image sequence showing that more time is required for trans-FRC migration in GlcNAc6ST-1 KO mice than in wild-type mice. The dotted lines indicate the boundary of FRCs. These images are serial single Z-frames (XY plane) and XZ or YZ cross sections. Lu, lumen; Pa, parenchyma. Scale bars,10 μm.

Fig. S5.

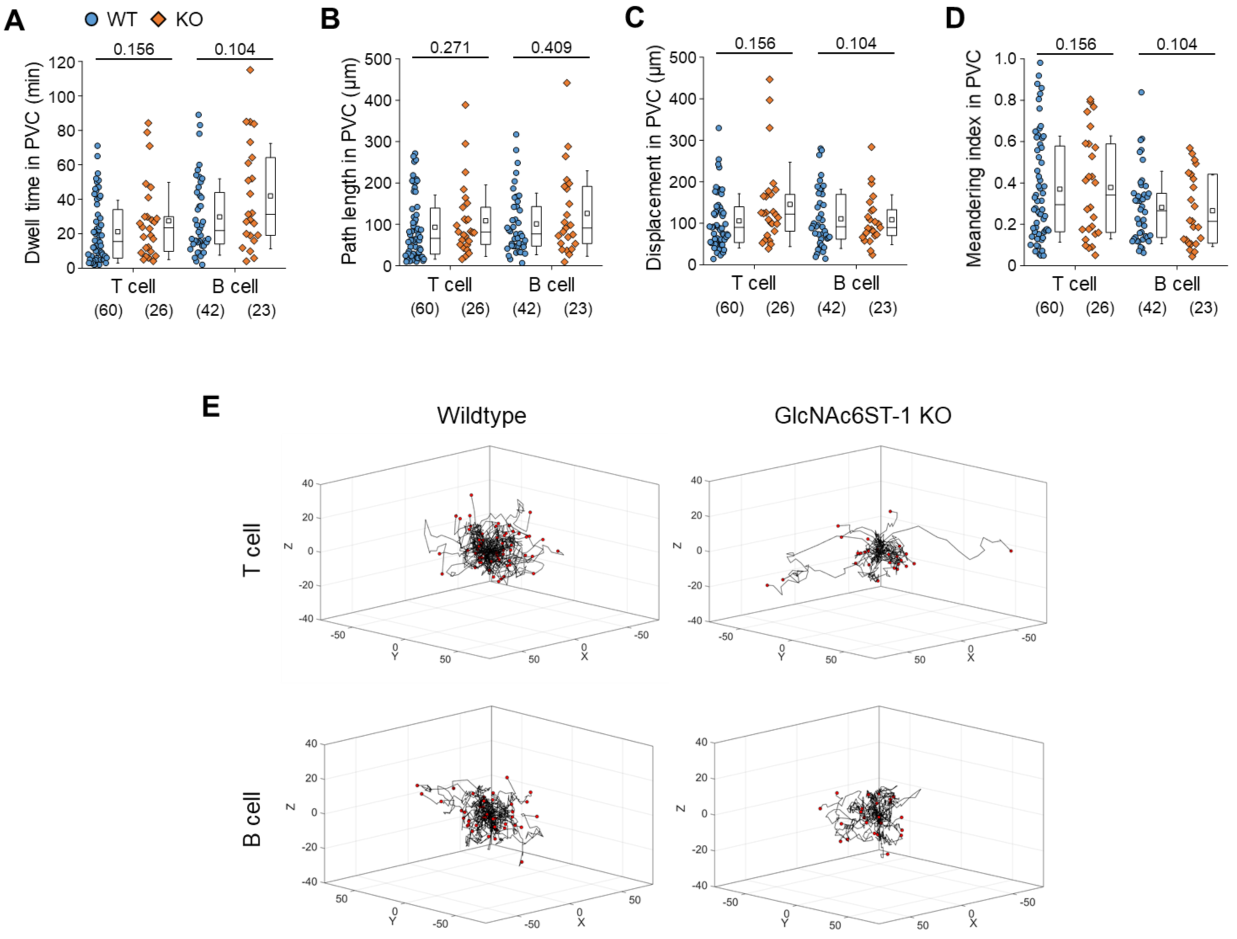


**Fig. S5. Effect of GlcNAc6ST-1 deficiency on T and B cell migration.**

(**A-D**) Quantitative analysis of T and B cell transmigration across HEV of GlcNAc6ST-1 KO mice compared with wildtype mice. There is no significant different between KO and WT mice in dwell time, path length, displacement and meandering index in PVC. The displacement in PVC indicates the straight distance from trans-EC migration site to trans-FRC migration site. The meandering index (MI) in PVC indicates the path length divided by the displacement. Each symbol represents one single. The box graph indicates 25$^{th}$ and 75$^{th}$ percentiles; the middle line and whiskers of the box indicate the median value and standard deviations, respectively; the small box represents the mean value. The number of analysed cells for 3 hours imaging is indicated below the graph. The overlapping T or B cells that were not distinguishable in PVC were excluded from analysis of intra-PVC migration. Four mice were analyzed for each group. P value was calculated with the Mann Whitney test. (**E**) 3D wind-rose plots show intra-PVC crawling paths of T or B cells as normalized for their trans-EC migration sites. The zero position in the coordinate and red dot of each track indicate the trans-EC migration site and trans-FRC migration site, respectively.

Fig. S6.

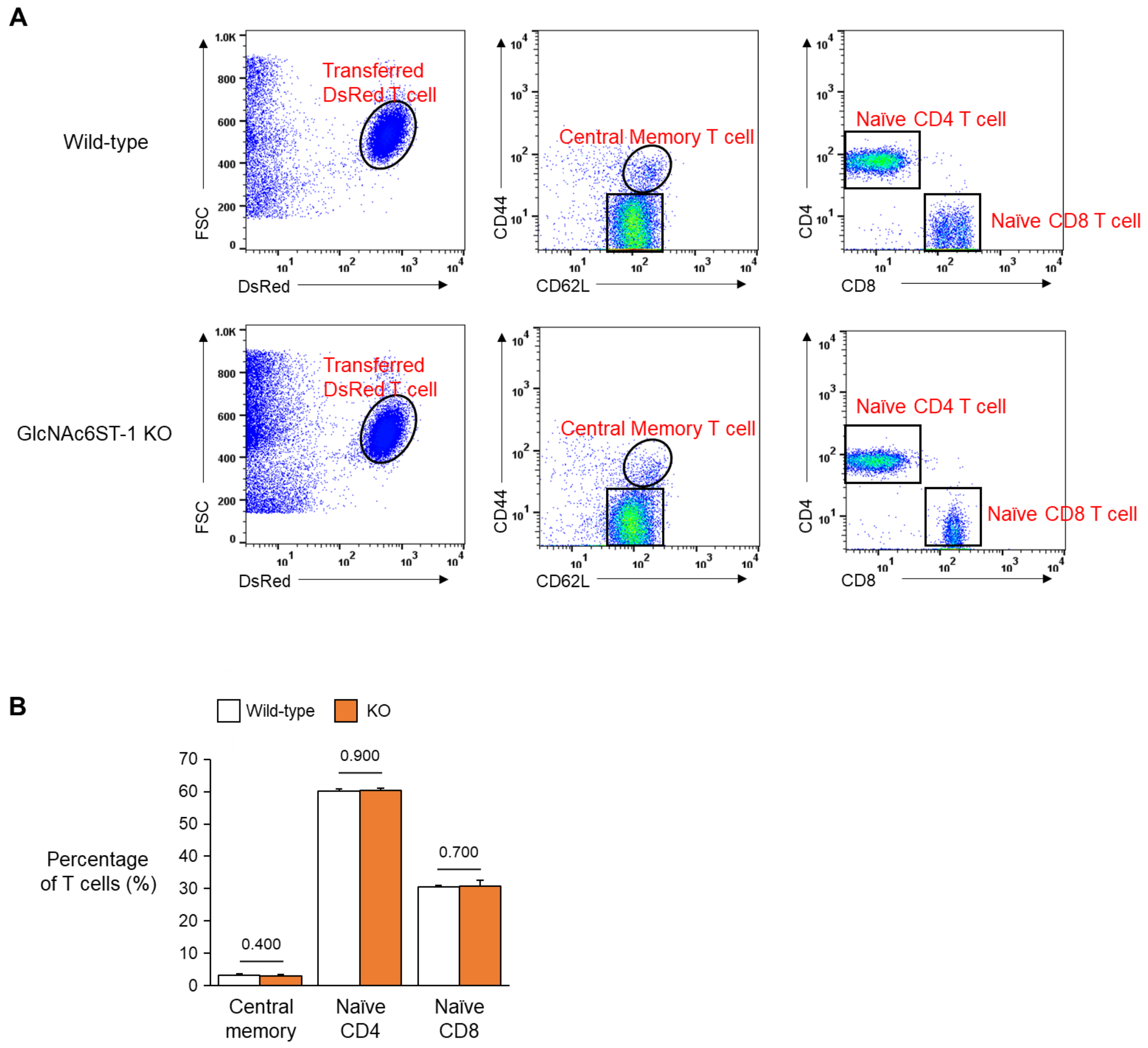


**Fig. S6. Percentage of homing T cell subsets between wild-type and GlcNAc6ST-1 KO mice.**

(**A**) We performed FACS analysis to investigate the percentage of DsRed T cell subsets (central memory, naïve CD4 and naïve CD8) recruited in peripheral lymph nodes (popliteal, inguinal) 3 hours after intravenous injection of DsRed pan-T cells into wild-type and GlcNAc6ST-1 KO mice. (**B**) No difference in the percentage of homing central memory, Naïve CD4 and CD8 T cells between wild-type and KO mice. Three mice were analysed for each group. P values were calculated with Mann-Whitney test.

**Fig. S7.**

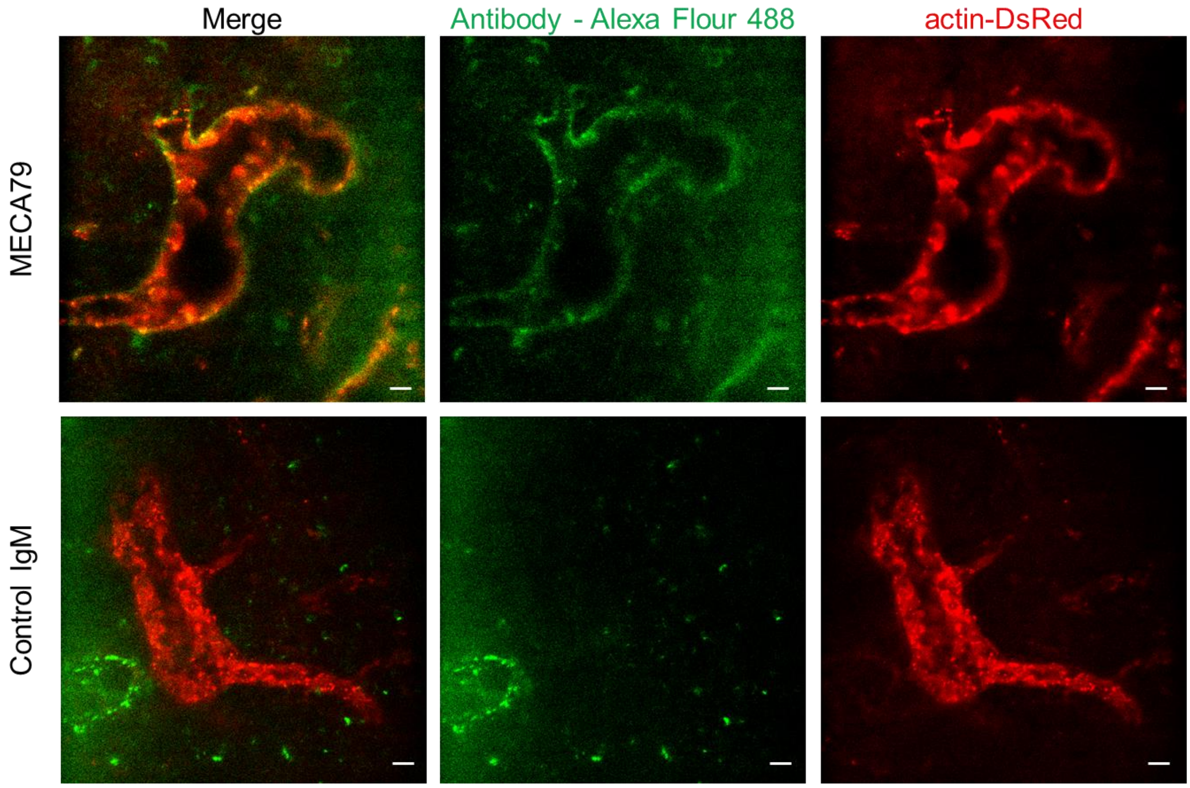


**Fig. S7. Fluorescence-labelled MECA79 distribution in HEV of lymph node after footpad injection.**

We injected MECA79 or Control IgM conjugated with Alexa Fluor 488 (10 μg, 20 μl) into a footpad of actin-DsRed mouse 3 hours before imaging. Actin-DsRed (red) shows an endothelium of HEV. MECA79 (green) highly accumulates in abluminal side of HEV while Control IgM is not shown in HEV. Scale bars, 10 μm.

**Fig. S8.**

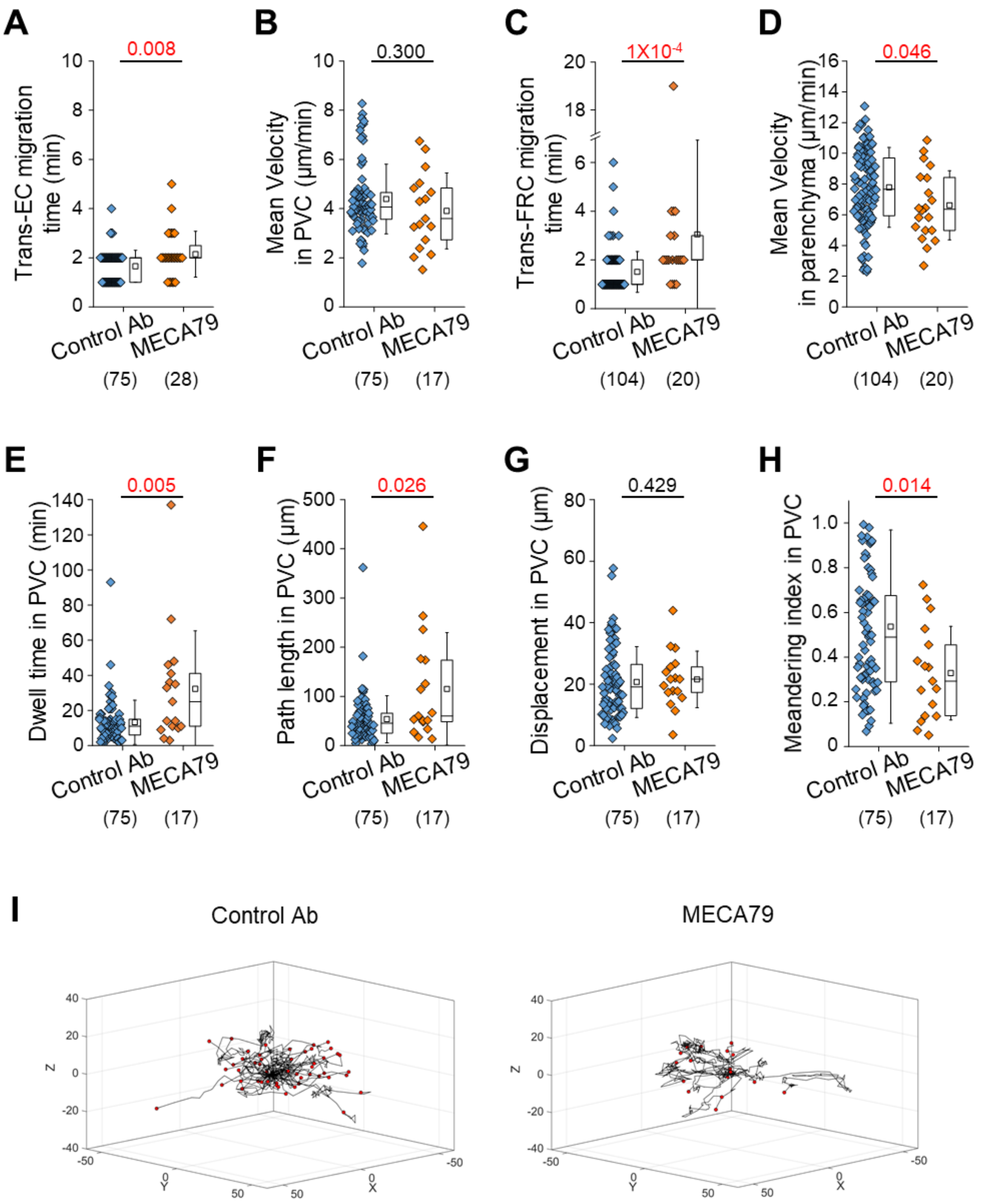


**Fig. S8. Effect of PNAd blockade on T cell transmigration across HEVs.**

MECA79 (a function-blocking antibody that reacts with the main L-selectin ligand PNAds) or control antibody was injected into a footpad 3 hours before imaging. (**A-H**)

Quantitative analysis of the migratory dynamics of the stepwise process of T cell transmigration across HEVs. The displacement in PVC indicates the straight distance from trans-EC migration site to trans-FRC migration site. The meandering index (MI) in PVC indicates the path length divided by the displacement. Each symbol represents a single cell. The box graph indicates the 25th and 75th percentiles; the middle line and whiskers of the box indicate the median value and standard deviation, respectively; the small square represents the mean value. The number of analysed cells is indicated below the graph. Five and 4 mice were analysed for the control Ab and MECA79 groups, respectively. P values were calculated with the Mann-Whitney test. (**I**) 3D wind-rose plots show intra-PVC crawling paths of T or B cells as normalized for their trans-EC migration sites. The zero position in the coordinate and red dot of each track indicate the trans-EC migration site and trans-FRC migration site, respectively.

Fig. S9.

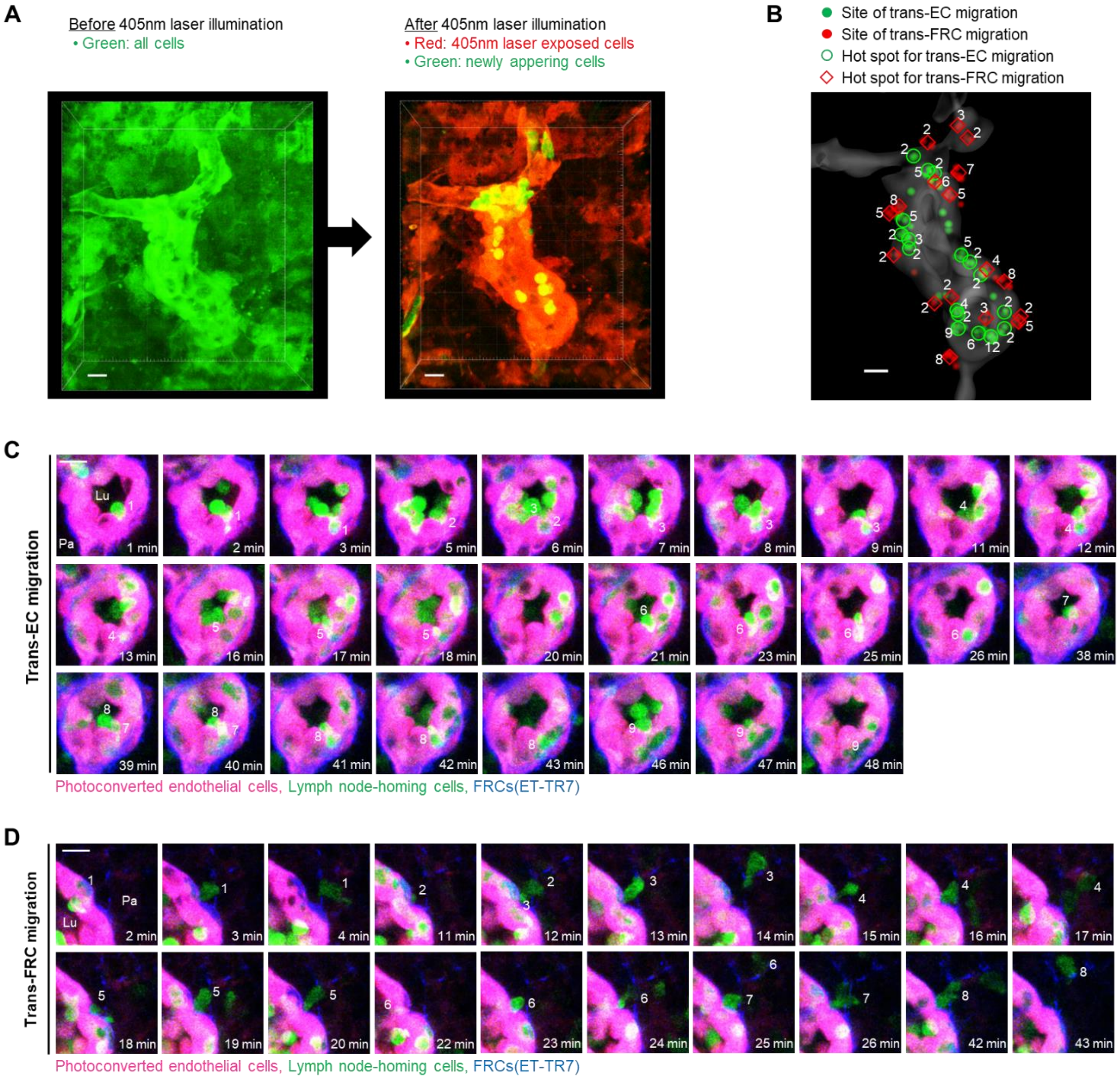


**Fig. S9. Hot spots of trans-EC and trans-FRC migrations for endogenous lymph node-homing cells.**

(**A**) Photoconversion of Kaede proteins in all cells in a field of view by 405 nm laser illumination. The laser exposed cells (photoconverted) change their color from green to red while newly appeared cells (nonphotoconverted) remain green. Photoconverted endothelial cells in HEV (red) can be clearly visualized because the endothelial cells express Kaede proteins at a level high enough to distinguish from the surrounding stromal cells and lymphocytes. (**B**) Representative 3D distribution of trans-EC (green dots) and trans-FRC migrations sites (red dots) of nonphotoconverted cells in the HEV (gray). The number of the nonphotoconverted cells transmigrating at same site (hot spot) is indicated. (**C**) Representative serial images showing that 9 nonphotoconverted cells (green) transmigrate across ECs (red) at same site. (**D**) Eight nonphotoconverted cells (green) transmigrate across FRCs (blue) at same site. Lu, lumen; Pa, parenchyma. These images (**C-D**) correspond to a 6 μm-thick maximum intensity projection. Scale bars,10 μm.

**Fig. S10.**

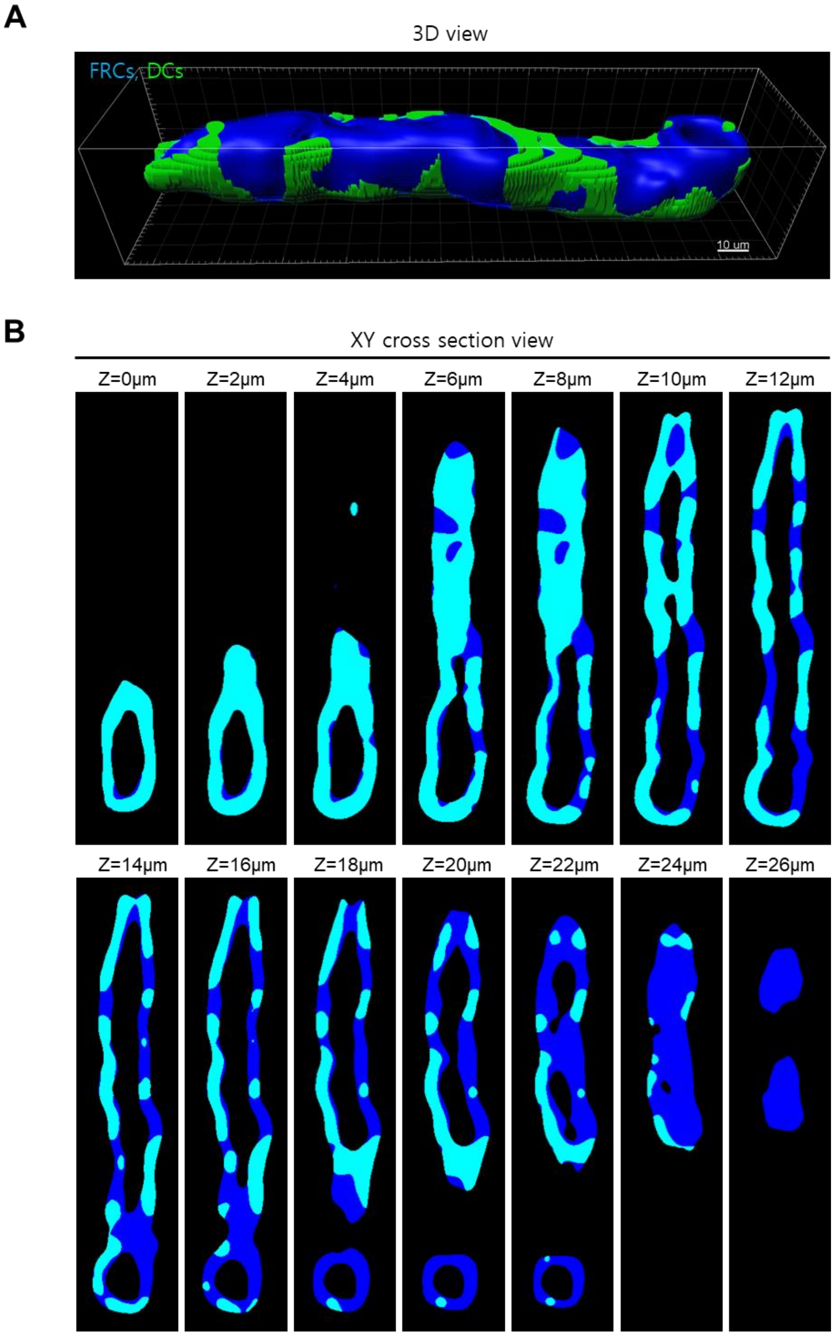


**Fig. S10.** Representative (**A**) 3D and (**B**) z-stack 2D images show DC coverage on HEV. We calculated the DC coverage on HEV by dividing FRC volume colocalized with DCs (cyan) by total FRC volume of HEV (blue). Scale bar, 10 μm.

**Fig. S11.**

■ T or B cell contact with DC during trans-FRC migration

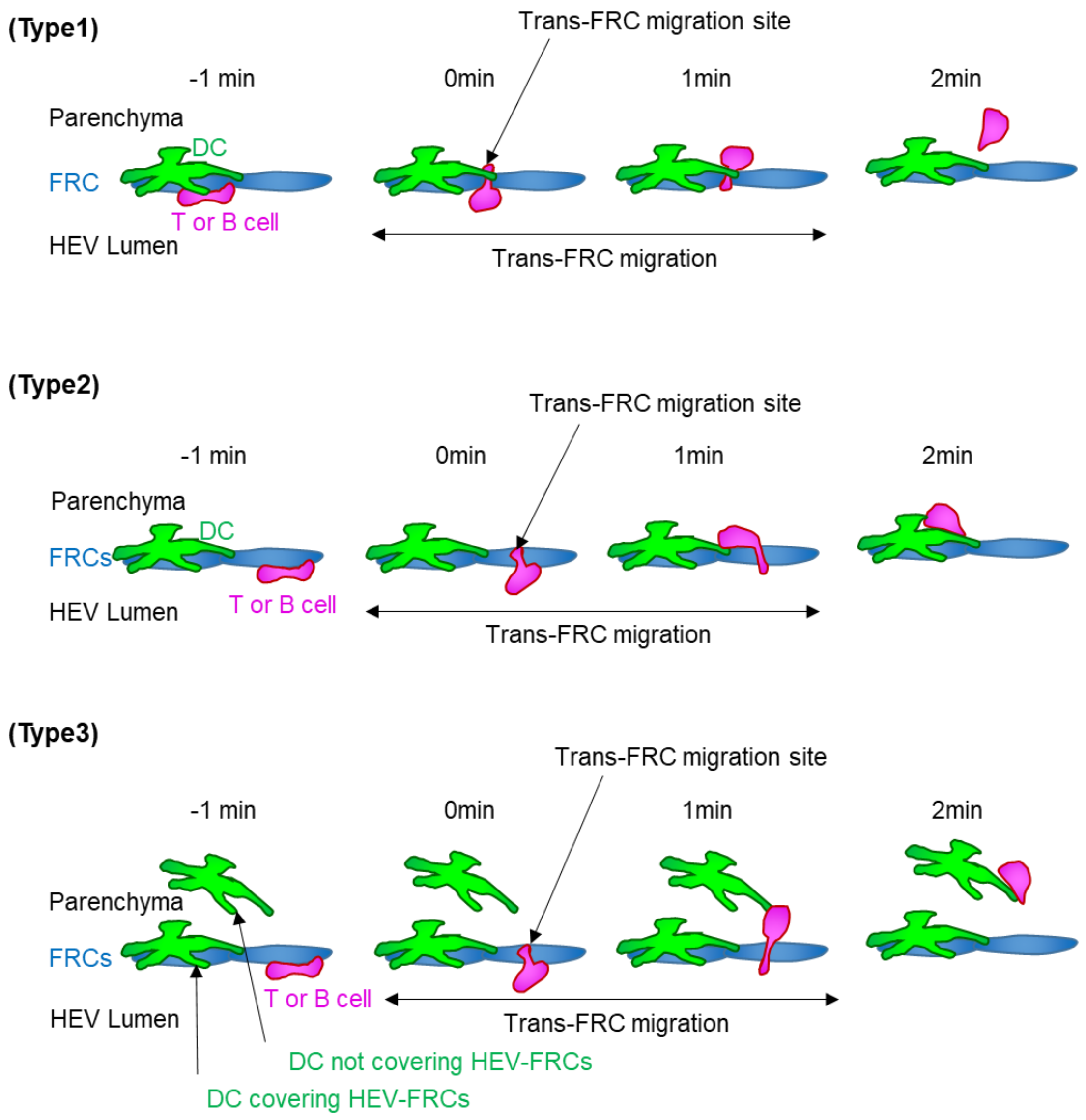


■ Trans-FRC migration without any contact with DCs

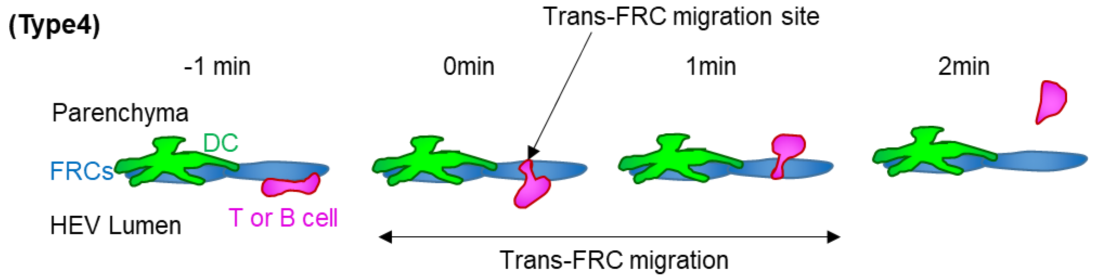

**Fig. S11. Classification of trans-FRC migration according to how T or B cell contacts with DCs.**

(**Type1**) 69 ± 10 % T cells and 66 ± 14 % B cells contact with DC covering their trans-FRC migration site during the trans-FRC migration. Type1 corresponds to Fig. 5D. (**Type2**) T or B cell contacts with DC covering HEV-FRCs although its trans-FRC migration site is not covered by the DC. (**Type3**) T or B cell contacts with DC not covering HEV-FRCs. 14 ± 9 % T cells and 22 ± 16 % B cells are of type 2 and type 3. (**Type4**) 18 ± 9 % T cells and 12 ± 9 % B cells do not contact with DCs during the trans-FRC migration. Mean ± SD. Eight (48 ± 18 T cells/mouse) and 8 mice (21 ± 19 B cells/mouse) were used for the analysis of T and B cells, respectively.

Fig. S12.

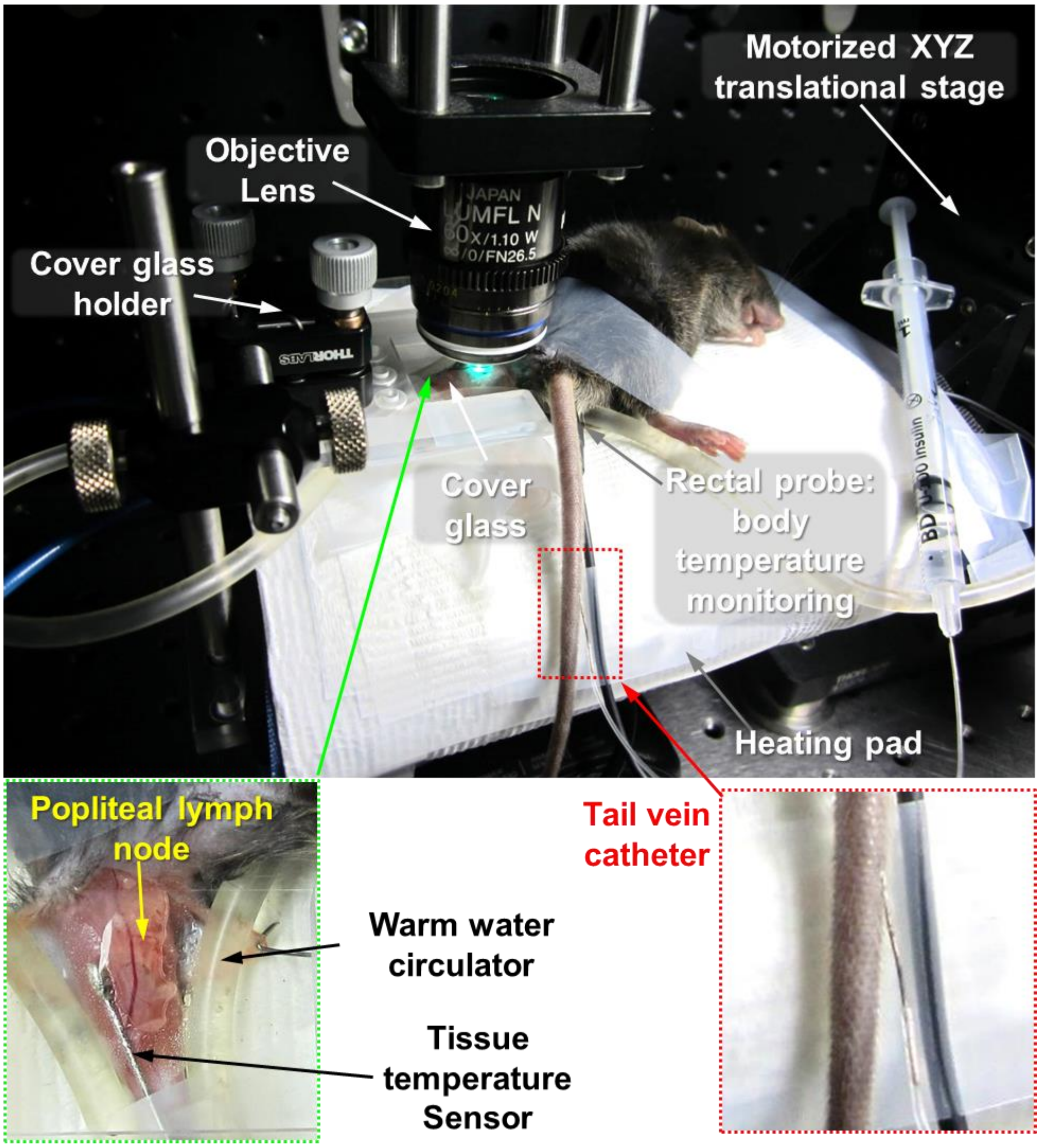


**Fig. S12. Mouse preparation for intravital imaging of a popliteal lymph node.**

To maintain a mouse body temperature, the mouse is placed on a heating pad where temperature is automatically controlled by receiving the mouse body temperature feedback with a rectal probe. Temperature of the exposed lymph node was maintained by warm water circulator and a tissue temperature sensor. A catheter is inserted into the tail vein of the mouse for injecting T and B cells immediately before the imaging and for injecting fluorescent dyes repeatedly during the imaging.

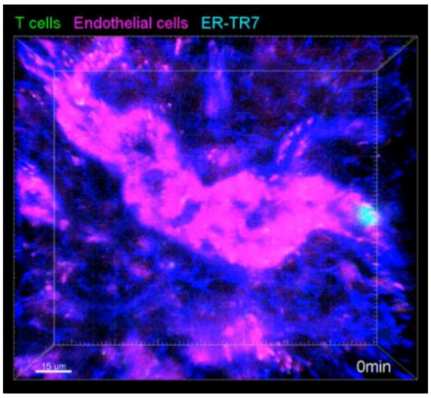


**Movie S1. Intravital 3D imaging of T cell transmigration across HEV.**

T cells (green), endothelial cells (red), ER-TR7 (blue). Scale bar, 15 μm.

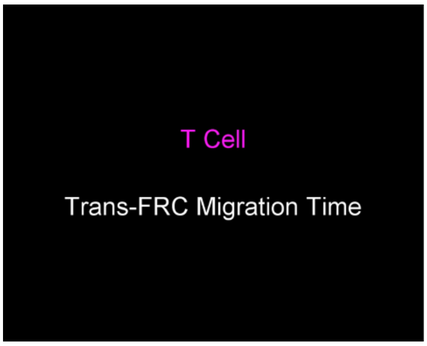


**Movie S2. Trans-FRC migration of T and B cells in wild-type and GlcNAc6ST-1 KO mice.**

GlcNAc6ST-1 deficiency leads to longer trans-FRC migration time for T and B cells. T cells (red), B cells (bright green), ER-TR7 (blue), HEV lumen (light green).

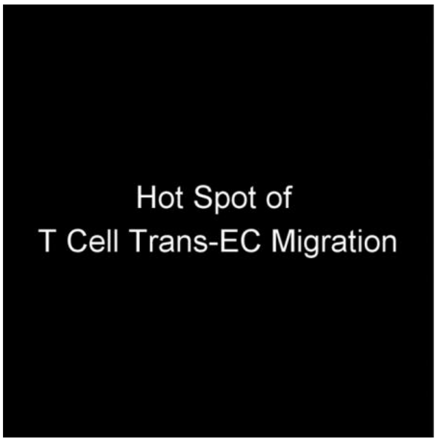


**Movie S3. Hot spot of T cell trans-EC migration.**

T cells (green), endothelial cells (red), ER-TR7 (blue).

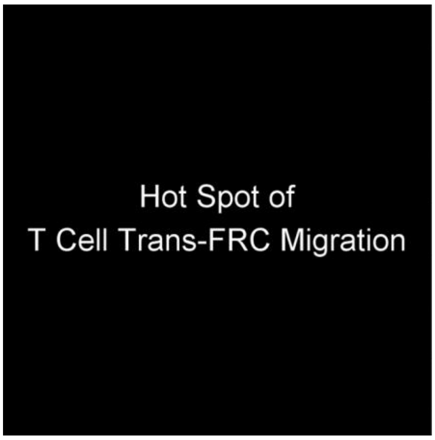


**Movie S4. Hot spot of T cell trans-FRC migration.**

T cells (green), endothelial cells (red), ER-TR7 (blue).

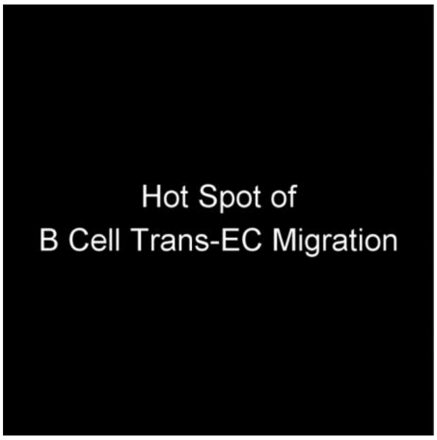


**Movie S5. Hot spot of B cell trans-EC migration.**

B cells (green), endothelial cells (red), ER-TR7 (blue).

Scale bar, 10 μm.

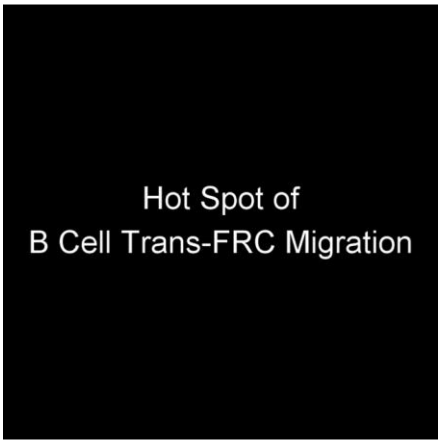


**Movie S6. Hot spot of B cell trans-FRC migration.**

B cells (green), endothelial cells (red), ER-TR7 (blue).

Scale bar, 10 μm.

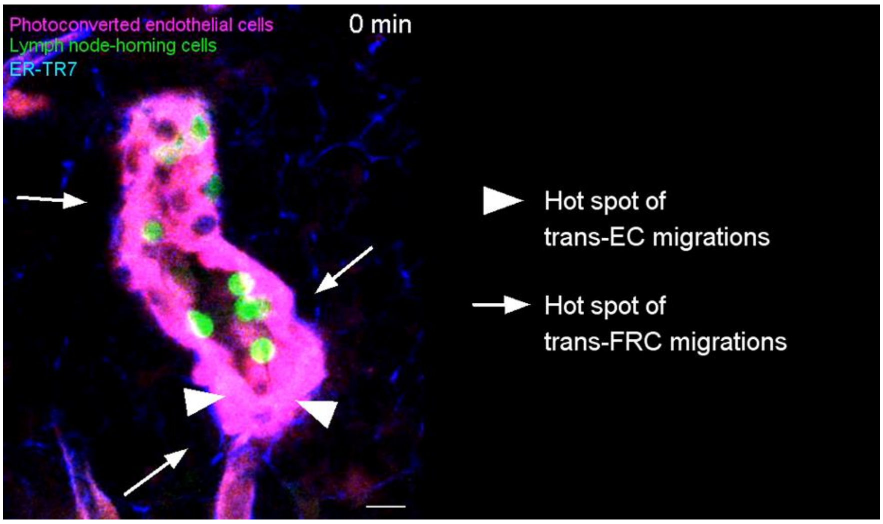


**Movie S7. Hot spot of trans-EC and trans-FRC migrations for endogenous lymph node-homing cells.** Photoconverted endothelial cells (red), lymph node-homing cells (green, non-photoconverted cells), ER-TR7(blue). The serial images correspond to a 6μm-thick maximum intensity projection. Scale bar, 10 μm.

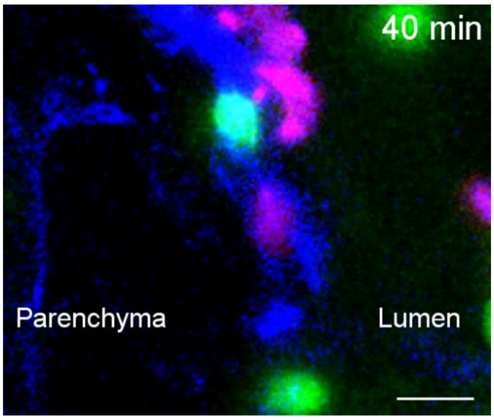


**Movie S8. T and B cells share the hot points of trans-FRC migration.**

T cells (red), B cells (bright green), ER-TR7 (blue), HEV lumen (light green). The serial images correspond to a 20μm-thick maximum intensity projection. Scale bar, 10 μm.

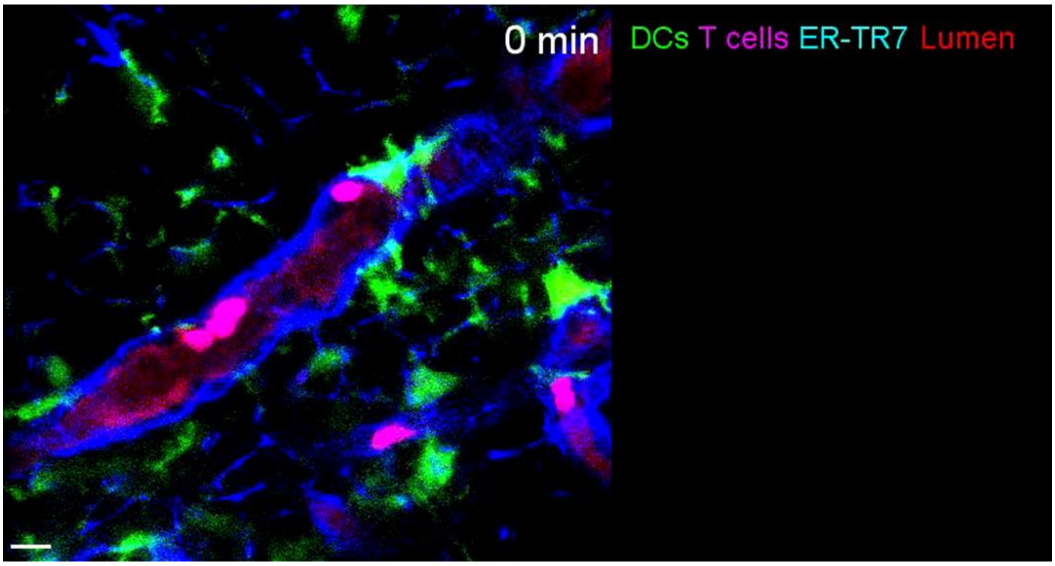


**Movie S9. T cells transmigrate across FRCs covered by CD11c+ DCs.**

DCs (green), FRCs (blue), T cells (bright red), HEV lumen (light red), Scale bar,10 μm. The serial images correspond to a 6 μm-thick maximum intensity projection. Scale bar,10 μm.
